# Brain Connectivity Patterns in Children Linked to Neurocognitive Abilities

**DOI:** 10.1101/2020.09.10.291500

**Authors:** Chandra Sripada, Mike Angstadt, Saige Rutherford, Aman Taxali, D. Angus Clark, Tristan Greathouse, Alex Weigard, Luke Hyde, Mary Heitzeg

**Author notes:** Correspondence: Chandra Sripada, Department of Psychiatry, University of Michigan, 4250 Plymouth Road, Ann Arbor MI 48109.

## Abstract

The development of objective brain-based measures of individual differences in psychological traits is a longstanding goal of clinical neuroscience. Here we show that reliable objective markers of children’s neurocognitive abilities can be built from measures of brain connectivity. The sample consists of 5,937 9- and 10-year-olds in the Adolescent Brain Cognitive Development multi-site study with high-quality functional connectomes that capture brain-wide connectivity. Using multivariate methods, we built predictive neuromarkers for a general factor of neurocognitive ability as well as for a number of specific cognitive abilities (e.g., spatial reasoning, working memory). Neuromarkers for the general neurocognitive factor successfully predicted scores for held-out participants at 19 out of 19 held-out sites, explaining over 14% of the variance in their scores. Neuromarkers for specific neurocognitive abilities also exhibited statistically reliable generalization to new participants. This study provides the strongest evidence to date that objective quantification of psychological traits is possible with functional neuroimaging.

## INTRODUCTION

Psychological traits and abilities arise from complex patterns in the structure and function of the human brain. A central goal for clinical neuroscience is to objectively measure these brain patterns in order to assess and predict individual differences in traits and abilities. Recent studies provide hints that objective quantification of psychological traits is possible with non-invasive functional neuroimaging^1–5^ (see ^6^ for a review), but modest sample sizes and inconsistent results have prevented any strong conclusions.

In the present work, we take a significant step forward. We establish the strongest evidence to date for the effectiveness and generalizability of brain-based objective markers (“neuromarkers”) for neurocognitive abilities, a set of inter-related abilities for reasoning, problem solving, manipulating representations, and learning and recall of information^7–9^. Individual differences in these abilities are important because they are associated with diverse life outcomes. Better neurocognitive abilities are associated with health, well-being, and occupational success^10,11^, while deficits in neurocognition, and closely related constructs such as executive functioning^7,9^, are associated with a broad range of psychopathologies^12–16^.

Traditionally, neurocognitive abilities were studied in functional imaging with task-based studies and locationist methodology^17–20^—participants are given tasks that engage neurocognitive processing with the aim of localizing task-associated processing to specific brain regions. Our work here differs in three respects. First, we take a network neuroscience approach^21,22^, examining interconnections among distributed large-scale networks measured when participants are at rest^23^. Second, we apply recently developed multivariate predictive modeling methods that aggregate (typically small) units of information across the entire brain, creating an overall best prediction of scores on individual difference variables.^24^ Such methods provide a substantial improvement in effect size compared to traditional mass univariate approaches (which conduct separate statistical tests at each brain feature), potentially allowing clinically meaningful predictions of individual differences at the level of the single subject. Third, we do not study how brain features are associated with a single neurocognitive task. Rather, we examine how brain connectivity is linked to an overarching general factor of neurocognitive ability^25–28^ that contributes to performance across diverse neurocognitive tasks, as well as several specific factors of neurocognitive ability^29,30^.

Our study leverages imaging and behavioral data from 11,875 9 and 10-year olds in the Adolescent Brain and Cognitive Development (ABCD) national consortium study, Release 2.1^31,32^. We applied bifactor modeling to the comprehensive ABCD 11-task neurocognitive battery^33^ to quantify individual differences in a dominant general factor of neurocognitive ability, which captured 75% of the variation in task scores (coefficient ω hierarchical^34^), as well as three domain-specific factors that together accounted for 13% of the variation in task scores (Figure 1). We also produced resting state connectomes for 5,937 youth who met stringent neuroimaging quality control standards; these connectomes capture tens of thousands of functional connections between hundreds of brain regions. We next applied a multivariate approach predictive modeling approach, brain basis set (BBS)^35,5,36^, to build neuromarkers for neurocognitive abilities from whole-brain functional connectivity patterns. To guard against identifying spurious relationships and to provide evidence of generalizability, we coupled BBS with leave-one-site-out cross-validation, in which we construct neuromarkers in all sites except one, test the marker at the held-out site, and repeat until each site is held out.

**Figure 1:**
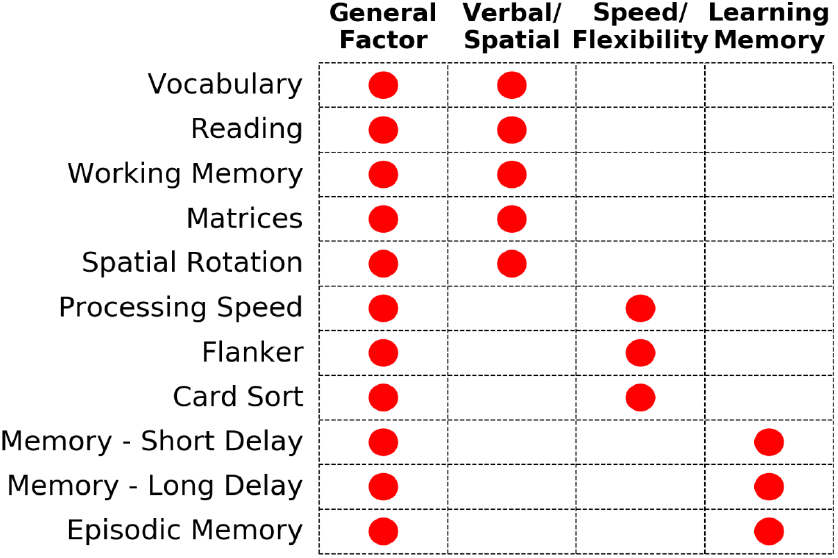
Factor Model of the ABCD Neurocognitive Battery. The comprehensive 11-task ABCD neurocognitive battery was factor analyzed yielding a general factor of neurocognitive ability and three specific factors.

**Figure 2:**
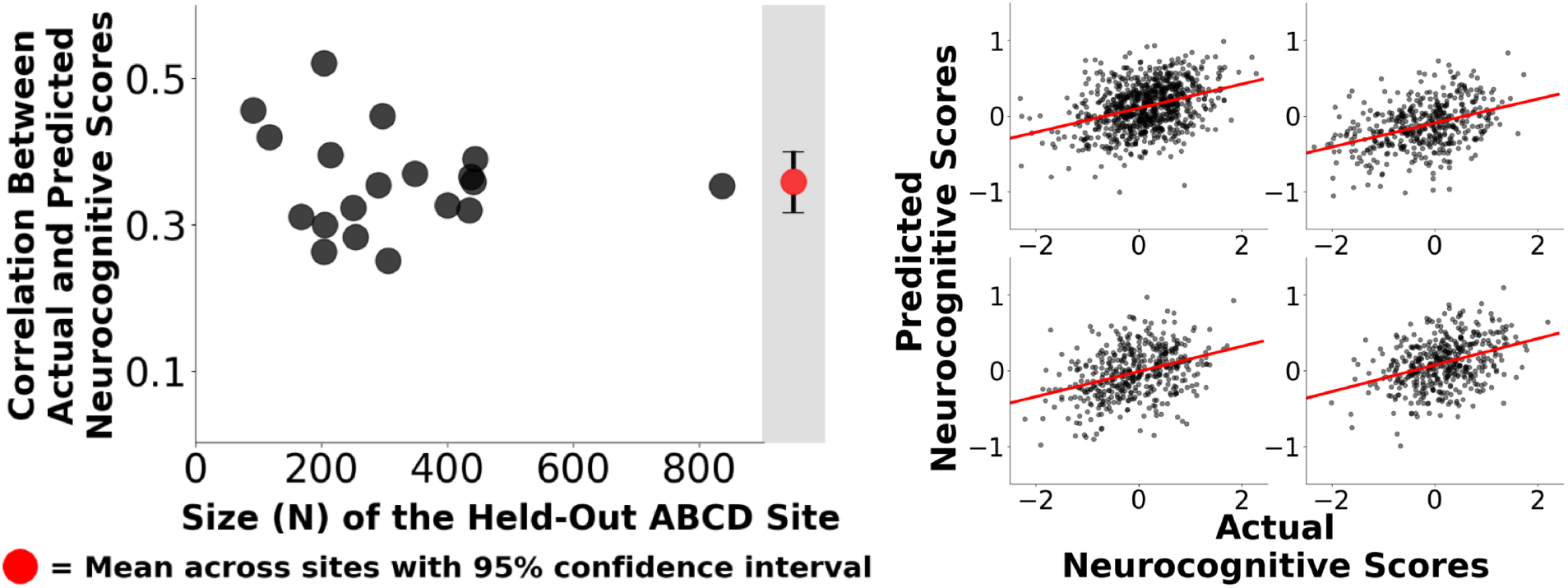
Correlations Between Predicted and Actual General Factor Scores in Leave-One-Site-Out Cross-Validation Analysis. Neuromarkers for the general factor of neurocognitive ability were constructed based on whole brain connectivity patterns. (Left Panel) These neuromarkers generalized to 19 out of 19 held out sites. The overall mean correlation between predicted and observed general factor scores for 5,937 held out subjects was 0.36, p_PERM_<0.0001 (observed correlation was higher than 10,000 correlations in the permutation distribution). (Right Panel) Scatter plots for the four largest held-out sites show highly consistent performance.

## RESULTS

### Connectivity-based neuromarkers for the general factor of neurocognitive ability are highly effective in predicting scores in held-out subjects

In leave-one-site-out cross-validation, the correlation between actual versus predicted general factor scores, averaging across folds of the cross-validation, was 0.36 (Figure 1, left panel). That is, brain connectivity patterns accounted for 14.2% of the variance in general factor scores in held-out samples of youth (variance explained was calculated with the r-squared cross-validated metric^37^). Cross-site generalizability was remarkably consistent (Figure 1, right panel). Correlations between predicted and actual scores were statistically significant in 19 out of 19 held-out sites (all 19 site-specific *p* values < 0.0001; observed correlations were higher than all 10,000 correlations in the permutation distribution).

### Predictive performance of neuromarkers for the general factor of neurocognition remained highly statistically significant across multiple robustness checks

We next assessed the robustness of our leave-one-site-out cross-validation analysis via sensitivity checks that modified key elements of the analysis stream. We tested alternative analyses that utilized: 1) Additional ABCD demographic covariates (household income, highest parental education, household marital status), and 2) A subsample restricted to participants who reported their race as White/European-American (non-Hispanic), to confirm that findings were not driven by potentially confounding demographic factors; 3) Bifactor models of neurocognition learned in the training sample and applied to the held-out sample (to create total separation between training and test samples); and 4) An ultra-low head motion sample (mean framewise displacement<0.2; to confirm motion was not responsible for any associations). As shown in Table 1, all results remained highly statistically significant across these changes, confirming the robustness of our analysis stream.

**Table 1:** Summary of Additional Analyses to Assess Robustness. The sensitivity of our main analyses (top row) to modeling choices was assessed with a number of robustness checks (rows 2-5). Correlations between actual and predicted neurocognitive scores remained statistically significant and similar in size across all these analyses. * = observed correlation was higher than all 10,000 correlations in the permutation distribution.

| Analysis | General Factor of Neurocognitive Ability | Number of Participants (N) |
| --- | --- | --- |
| 1. Main Analysis | $r=0.36$; $p<0.0001^*$ | 5,937 |
| 2. Additional Demographic Controls | $r=0.30$; $p<0.0001^*$ | 5,468 |
| 3. White/European-American (non-Hispanic) Subsample | $r=0.27$; $p<0.0001^*$ | 3,480 |
| 4. Leave-One-Site Out General Factor | $r=0.36$; $p<0.0001^*$ | 5,937 |
| 5. Ultra-Low Motion Subsample | $r=0.32$; $p<0.0001^*$ | 2,847 |

### Neuromarkers for specific cognitive abilities also exhibited statistically reliable generalization to unseen subjects, with neuromarkers for verbal abilities performing the best

We next assessed the predictivity of neuromarkers for specific neurocognitive abilities, in particular the three specific factors as well as the 11 individual neurocognitive tasks (Figure 1). Neuromarkers were constructed in 14 separate BBS models, each tested with leave-one-site-out cross-validation. We found that neuromarkers for all 14 specific ability variables exhibited statistically reliable generalization to unseen subjects (Figure 3). Neuromarkers for verbal abilities (vocabulary and reading) performed the best, each accounting for more than 9% of the variance in scores in held out participants. To address the fact that the general factor is correlated (to varying degrees) with some of the 14 specific ability variables, we constructed additional neuromarkers for all 14 specific ability variables, this time performing leave-one-site-out cross-validation controlling for the effect of the general factor of neurocognitive ability. We found 13 of the 14 neuromarkers continued to exhibit statistically reliable generalization to unseen subjects (the 14^th^ neuromarker, working memory, was trend significant at *p*_PERM_=0.10, Figure S3). This result demonstrates that resting state connectivity patterns contain unique information about multiple specific domains of neurocognitive abilities over and above information about the general factor.

**Figure 3:**
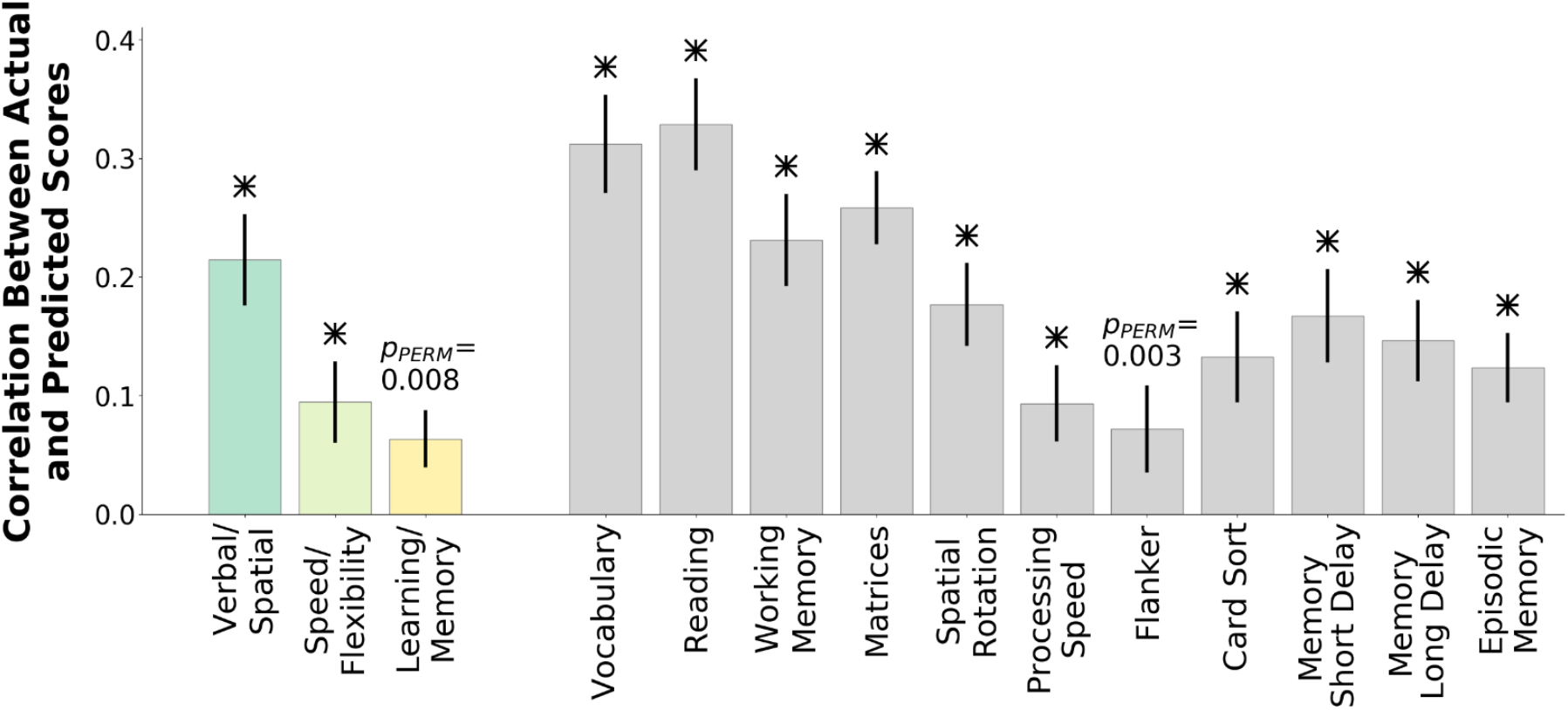
Correlations Between Predicted and Actual Scores for 14 Specific Neurocognitive Abilities in Leave-One-Site-Out Cross-Validation Analysis. Neuromarkers for 14 specific ability variables were constructed based on whole brain connectivity patterns. All 14 neuromarkers exhibited statistically reliable generalization to unseen subjects.

### Functional connections involving control networks and processing networks were prominent in the general factor neuromarker. Additionally, these connections varied the most across the fifteen neuromarkers for different neurocognitive abilities

We next examined consensus connectomes, importance-weighted composite maps associated each neuromarker (see Methods, §9). For the general factor neuromarker, its consensus connectome (Figure 4) showed prominent representation of connections involving control networks (fronto-parietal, cingulo-opercular, ventral attention, and dorsal attention) and processing networks (visual, default). Connections involving these networks were 56.4% of the suprathreshold connections in the general factor neuromarker, even though make up only 22.6% of the connections in the connectome.

**Figure 4:**
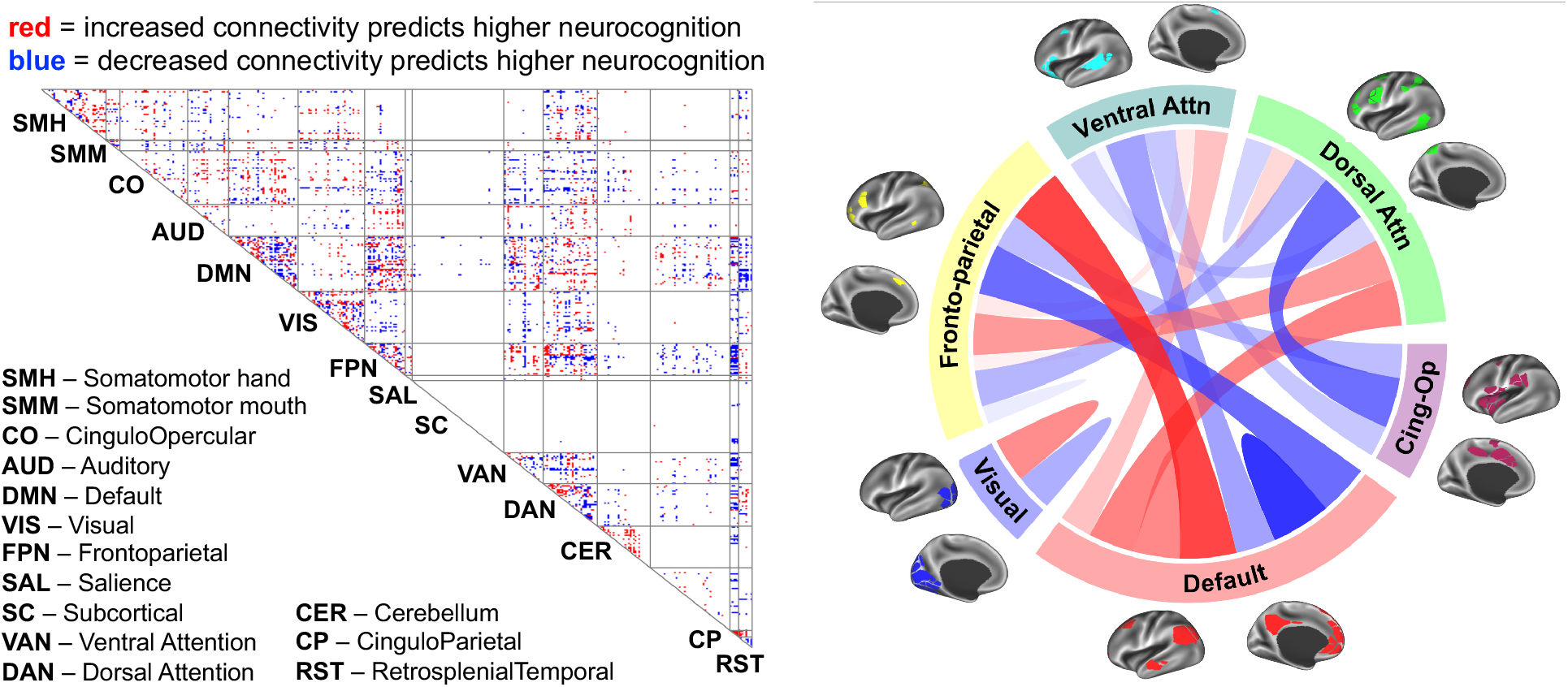
Connections Between Brain Networks in the Neuromarker for the General Factor of Neurocognition. (Left Panel) Supra-threshold connections linking large-scale networks are shown as red and blue dots. The map shows a distributed set of brain-wide connections is related to the general factor for neurocognitive ability. Connections involving control networks (fronto-parietal, ventral attention, dorsal attention, and cingulo-opercular) and processing networks (visual, default) are especially prominent. (Right Panel) Network map highlighting connections involving control networks and processing networks. Width of chords represents number of suprathreshold connections linking the indicated pair of networks. Red shades = connections at which higher connectivity predicts higher neurocognitive scores; Blue shades = connections at which lower connectivity predicts higher neurocognitive scores.

We next calculated a variability index (Methods, §10) that identifies pairs of networks whose connections varied the most across the 15 neuromarkers for neurocognitive abilities (the index separates positive and negative internetwork connections, and is thus calculated over 240 inter-network values). This index illuminates the main network connectivity patterns that best differentiate the neuromarkers. We found the top 21 inter-network relationships (out of 240 total) accounted for over 50% of the variance in inter-network connectivity across the 15 neuromarkers. These 21 highly varying inter-network relationships are shown in Figure 5, which in addition highlights interesting patterns of specificity for different domains of neurocognition. For example, the Speed/Flexibility factor is heavily represented in negative connections involving fronto-parietal and cingulo-opercular connections with default network. The Learning/Memory factor, in contrast, is most represented in negative connections within default network and positive connections linking default network with fronto-parietal network. Similar patterns of specificity are found for other specific neurocognitive abilities. Additionally, we found connections linking control networks and processing networks (see Figure 5 caption), which were heavily represented in the general factor neurosignature (see Figure 4), were also the most variable across neuromarkers.

**Figure 5:**
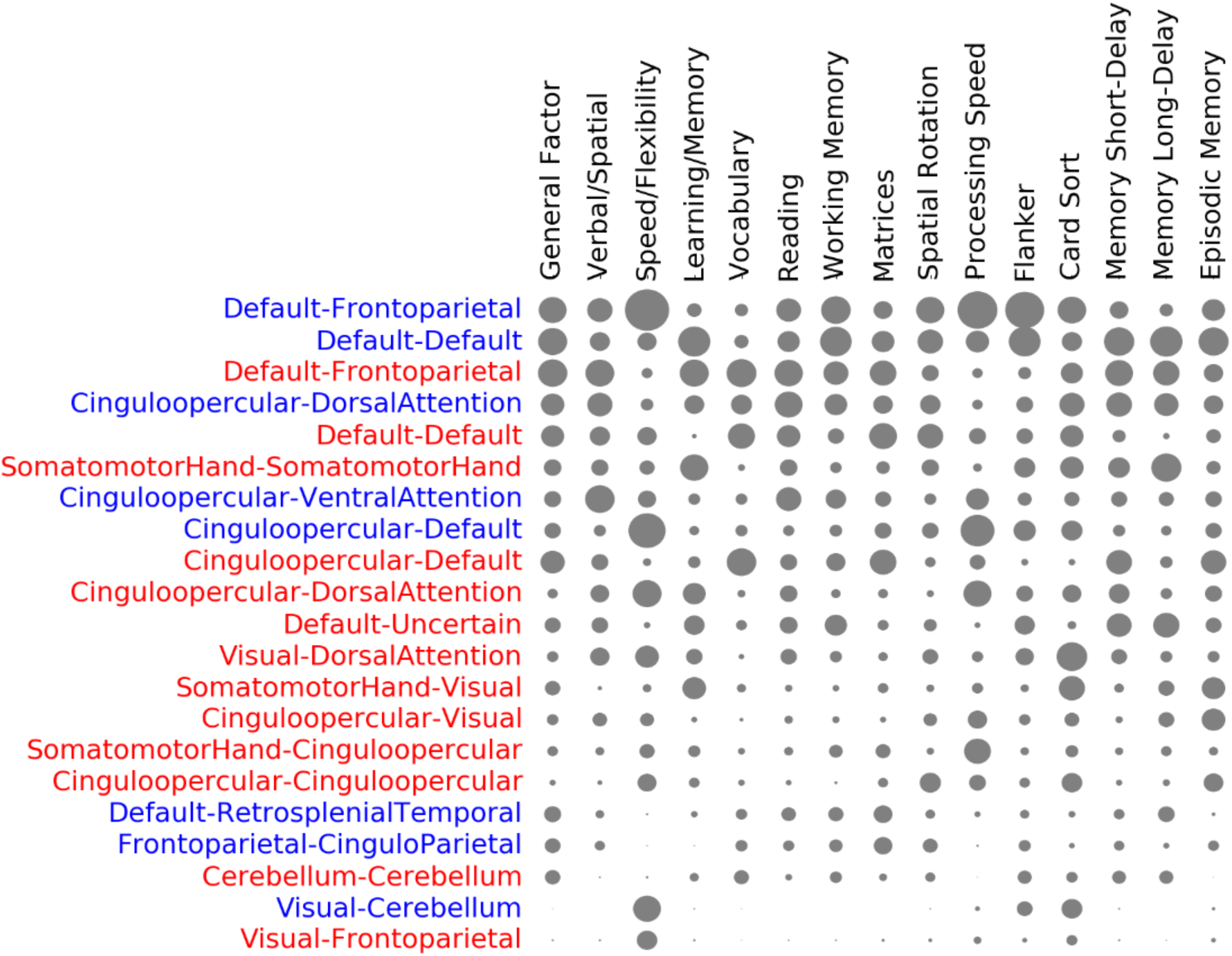
Connections Involving Control Networks and Processing Networks Differentiate Neuromarkers for Different Neurocognitive Abilities. We identified pairs of networks (listed in rows) in which inter-network connections varied the most across the fifteen neuromarkers for neurocognitive abilities (listed in columns). The size of the gray circles indicates the number of suprathreshold connections linking the pair of networks for each neuromarker. Red network labels indicate positive connections and blue indicate negative connections. The figure shows that connections involving control networks (Fronto-parietal, Cingulo-opercular, Dorsal Attention, Ventral Attention) and processing networks (Default, Visual, Somatomotor-Hand) play a major role in differentiating neuromarkers for distinct neurocognitive abilities.

## DISCUSSION

This study examined behavioral and resting state imaging data for 5,937 9- and 10-year-old participants across 19 sites in the ABCD Consortium study^33,38^. Using a multivariate predictive modeling approach, we aimed to build neuromarkers for neurocognitive abilities from resting state connectivity patterns and validate their performance with leave-one-site cross-validation. Our main findings are that multivariate neuromarkers of neurocognitive abilities are effective at predicting multiple domains of neurocognition and reliably generalize to held out subjects. In addition, neuromarkers for the general factor of neurocognition were particularly effective, explaining over 14% of variance in scores in held out subjects, a clinically meaningful level of predictivity, and remained highly predictive after a number of robustness checks. Our results provide the strongest evidence yet that neuroimaging can be used to build reliable and generalizable brain-based markers for psychological traits and abilities. Moreover, these findings set the stage for additional investigation in the longitudinal ABCD dataset to better understand how the connectivity patterns here linked to neurocognitive abilities change and mature over the course of adolescence.

Previous studies also identified links between brain imaging features, including resting state functional connectivity, and neurocognitive phenotypes^1–6^. The present study adds to the literature in three key ways. First, this is the largest study ever examining links between brain connectivity and cognitive abilities in youth. Larger samples enable estimation of effects with less variability^39^, yielding especially reproducible insights into brain-behavior relationships. Second, this study provides unique evidence about generalizability, showing that neuromarkers trained in one set of subjects effectively and consistently generalize to new subjects (with successful generalization in 19 out of 19 held out ABCD sites). The utility of imaging-based markers for psychological traits and abilities depends heavily on their applicability to new datasets collected at heterogenous sites with different subject characteristics and scanners, and this study confirms that strong generalizability is possible. Third, most previous neuroimaging studies exclusively examined a single aspect of cognition, such as a single neurocognitive ability^1,4,40^, or a general factor of cognitive ability^3,36,41^. This study is among the first to shed light on how brain connectivity contributes to the general factor of neurocognition, several specific factors, as well as a number of individual neurocognitive abilities.

To this point, we found that neuromarkers across different domains of neurocognition exhibited varying levels of performance. Predictivity was highest for the general factor of neurocognitive ability, explaining over 14% of the variance in scores in out of sample subjects. Strong predictivity was also observed for a verbal/spatial specific factor as well as for several individual tasks, especially for the crystalized abilities of reading and vocabulary. Relatively poorer performance, on the other hand, was observed for two other neurocognitive domains, the Speed/Flexibility domain and the Learning/Memory domain, as well as the individual task associated with these domains (Figure 3).

In explaining these differences, it is possible that these neurocognitive variables that were less well predicted simply do not have sizable signatures in resting state connectomes in 9- and 10-year old youth. There is some evidence that neurocognitive abilities remain fairly undifferentiated^27^, unspecialized^42^, or underdeveloped^43,44^ in late childhood/early adolescence. Thus, it is possible that performance for a number of specific factors and individual tasks will improve as the architecture of neurocognition becomes more refined over the course of adolescence and the specific abilities reflected in these factors/tasks separate out from the general factor. An alternative explanation is that the multivariate classifiers used for the present analysis are not sensitive to signatures for certain specific factors and individual tasks. For example, if resting state signatures of certain neurocognitive abilities are mainly localized in small, spatially discrete structures (e.g., striatum or hippocampus), classifiers that rely on distributed whole-connectome information, including the BBS method used in the present report, will be unlikely to recover them. In future studies, alternative classifiers (e.g., supervised methods), modalities (e.g., task-based methods), or search strategies (e.g., regions of interest approaches) could be utilized.

The connectivity neurosignatures for the general factor of neurocognition and other neurocognitive variables were highly distributed across the connectome, but there was nonetheless a concentration of connections involving control networks (such as fronto-parietal network and cingulo-opercular network) and processing networks (such as default network and visual network). Additionally, we found control network/processing network connections were the most variable across the set of neurosignatures for different neurocognitive domains. Control networks^45,46^ are proposed to be the source of cognitive control signals that modulate responses in other networks based on contextual demands—a function that is consistent with the observation that these networks are implicated in higher-order cognitive abilities. Additionally, connections involving control networks and default network are found to be among the most variable across individuals.^47^ These connections also exhibit substantial maturation across childhood and adolescence^48,49^, and their maturational trajectories exhibit significant inter-individual differences^4,50,51^. Taken together, these observations align well with our finding that connections involving control networks and processing networks are a primary locus of individual differences in neurocognitive abilities.

One important potential application for neuromarkers of neurocognition is in elucidating psychological, neural, and developmental mechanisms of cognitive abilities^43,52^. Neuromarkers summarize complex distributed whole-brain connectivity patterns with a small number of quantitative metrics^24^, opening the door to sophisticated statistical modeling methods that assess genetic and environmental (e.g., poverty, family environment) contributions to neuromarker expression and, in turn, the contributions of neuromarker expression to subsequent outcomes. Critically, a major strength of the 10-year longitudinal ABCD study^31^ is that it allows researchers to track the maturational patterns across adolescence of the connections implicated in our neurocognitive markers. This will facilitate future work that delineates in detail the role of neuromarker expression in mediating the relationship between genetic and environmental risk factors (e.g., poverty, family environment) and the subsequent emergence of behavioral (e.g., substance initiation) or psychopathological (e.g., psychosis) endpoints.

Additionally, neuromarkers of the type demonstrated here could one day find more direct clinical application. There is growing evidence that impaired neurocognitive abilities are associated with diverse forms of psychopathology^12–16^, including schizophrenia^53–55^, externalizing disorders such as ADHD^56,57^ and substance use disorders^58,59^, and internalizing disorders such as depression^60,61^. Notably, in this same ABCD baseline sample, we recently showed^62^ that reduced scores on the general factor of neurocognition were associated with elevation in the general factor of psychopathology^63–65^ (widely terms the “P factor”), which confers vulnerability to nearly all prevalent psychiatric symptoms. The possibility thus exists that neuromarkers for neurocognitive abilities could find clinical use in identifying individuals at risk for negative psychiatric outcomes, potentially at an early age well before overt signs and symptoms have emerged.

In sum, in a large rigorously characterized sample of youth, we established that neuromarkers of neurocognitive abilities built from distributed brain network connectivity patterns consistently generalize to new subjects across different data collection sites, are robust across a number of sensitivity checks, and capture clinically meaningful quantities of variation in general and specific neurocognitive abilities.

## METHODS

### 1. Sample and Data

The ABCD study is a multisite longitudinal study with 11,875 children between 9-10 years of age from 21 sites across the United States. The study conforms to the rules and procedures of each site’s Institutional Review Board, and all participants provide informed consent (parents) or assent (children). Detailed description of recruitment procedures^66^, assessments^67^, and imaging protocols^38^ are available elsewhere.

### 2. Data Acquisition, fMRI Preprocessing, and Connectome Generation

Imaging protocols were harmonized across sites and scanners. High spatial (2.4 mm isotropic) and temporal resolution (TR=800 ms) resting state fMRI was acquired in four separate runs (5min per run, 20 minutes total, full details are described in ^68^). The entire data pipeline described below was run through automated scripts on the University of Michigan’s high-performance cluster, and is described below, with additional detailed methods automatically generated by fRMIPrep software provided in the Supplement. Code for running the analyses can be found at Code for running the analyses can be found at https://github.com/SripadaLab/ABCD_Resting_Neurocognition.

Preprocessing was performed using fMRIPrep version 1.5.0^69^, a Nipype^70^ based tool. Full details of the fMRIPrep analysis can be found in supplemental materials. Briefly, T1-weighted (T1w) and T2-weighted images were run through recon-all using FreeSurfer v6.0.1. T1w images were also spatially normalized nonlinearly to MNI152NLin6Asym space using ANTs 2.2.0. Each functional run was corrected for fieldmap distortions, rigidly coregistered to the T1, motion corrected, and normalized to standard space. ICA-AROMA was run to generate aggressive noise regressors. Anatomical CompCor was run and the top 5 principal components of both CSF and white matter were retained. Functional data were transformed to CIFTI space using HCP’s Connectome Workbench. All preprocessed data were visually inspected at two separate stages to ensure only high-quality data was included: After co-registration of the functional data to the structural data and after registration of the functional data to MNI template space.

Connectomes were generated for each functional run using the Gordon 333 parcel atlas^71^, augmented with parcels from high-resolution subcortical^72^ and cerebellar^73^ atlases. Volumes exceeding a framewise displacement threshold of 0.5mm were marked to be censored. Covariates were regressed out of the time series in a single step^74^, including: linear trend, 24 motion parameters (original translations/rotations + derivatives + quadratics), aCompCorr 5 CSF and 5 WM components and ICA-AROMA aggressive components, high pass filtering at 0.008Hz, and censored volumes. Next, correlation matrices were calculated for each run. Each matrix was then Fisher r-to-z transformed, and then averaged across runs for each subject yielding their final connectome.

### 3. Bifactor Modeling of the ABCD Neurocognition Task Battery

Preliminary exploratory factor analyses (EFA) were first conducted (maximum likelihood estimation with oblique geomin rotation) to explore the latent structure of the 11 ABCD neurocognitive tasks. In addition, a parallel analysis with 1,000 random draws was also run that suggested the presence of three factors. The optimal factor structure was determined by considering the scree plot, parallel analysis, model fit, and the interpretability of different factor solutions (including consistency with past research). Overall, three broad factors best characterize these tasks, corresponding to spatial/verbal, speed/flexibility, and learning/memory; in the bifactor models these three factors served as the specific factors. Follow-up confirmatory factor analysis showed very good fit by conventional standards (χ^2^ (34)=443.16, *p*<0.001, RMSEA=0.03, CLI=0.99, TLI=0.98, SRMR=0.02), with the general factor capturing 75% of the variation in task scores (coefficient ω hierarchical^34^), and the three domain-specific factors together accounting for 13% of the variation in task scores (see Figure S3).

### 4. Inclusion/Exclusion

There are 11,875 subjects in the ABCD Release 2.0.1 dataset. Screening was initially done using ABCD raw QC to limit to subjects with 2 or more good runs of resting data as well as a good T1 and T2 image (QC score, protocol compliance score, and complete all =1). This resulted in 9598 subjects with 2 or more runs that entered preprocessing. Each run was subsequently visually inspected for registration as well as for warping quality, and only those subjects who still had 2 or more good runs were retained (N=8858). After connectome generation, runs were excluded if they had less than 4 minutes of uncensored data, and next subjects were retained only if they had 2 or more good runs (N=6568). Next, sites with fewer than 75 subjects were dropped. This left us with N=6449 subjects across 19 sites to enter PCA. Finally, subjects with missing values for neurocognitive scores or nuisance covariates were excluded. This left 5937 subjects to enter our main BBS predictive modeling analysis. Number of subjects for alternative analysis streams conducted as part of our robustness checks are reported in Table 1.

### 5. Constructing a Brain Basis Set (BBS)

BBS is a validated multivariate method that uses principal components dimensionality reduction to produce a basis set of components that are then associated with phenotypes^4,35^. We select the top 250 components for our basis set based on previous work showing that 50-100 components per 1000 subjects captures most meaningful variance without overfitting^35,36^.

### 6. Leave-One-Site-Out Cross Validation

To assess generalizability of BBS-based regression models, we used leave-one-site-out cross-validation. In each fold of the cross-validation, data from one of the 19 sites served as the held-out test dataset and data from the other 18 sites served as the training dataset. Additionally, to ensure separation of train and test datasets, at each fold of the cross-validation, a new PCA was performed on the training dataset yielding a 250-component basis set. We assessed the performance of BBS models with Pearson’s correlation, cross-validated r-squared (see Supplement), and mean squared error.

### 7. Accounting for Covariates in Cross-Validation Framework

In each fold of cross-validation, BBS models were trained in the train partition with the following covariates: gender, race, age, age squared, mean FD and mean FD squared. To maintain strict separation between training and test datasets, regression coefficients for the covariates learned from the training sample are applied to the test sample, and the variance they explain is subtracted away. This procedure, described in detail in our previous publication^36^, yields an estimate of the contribution of brain components alone in predicting test subject P factor scores, excluding the contribution of the nuisance covariates. Note that by employing leave-one-site-out, members of twinships and sibships are never present in both training and test samples.

### 8. Permutation Testing

We assessed the significance of all cross-validation-based correlations with non-parametric permutation tests in which we randomly permuted the 5,880 subjects’ P factor scores 10,000 times, as described in detail in the Supplement.

### 9. Consensus Connectome Maps

To help convey overall patterns across entire BBS multiple regression models with 250 components, we constructed “consensus” component maps. We used multi-level multiple regression modeling, with the neurocognitive scores as outcome variables and expression scores for the 250 components as predictors. Gender, race, age, mean FD, and mean FD squared were entered as fixed effect covariates, with family id and ABCD site entered as random effects (family nested within site). We next multiplied each connectomic component with its associated regression coefficient. We then summed across all 250 components yielding a single map, and thresholded the entries at *z*=2. We in addition created circular visualizations of consensus connectomes (see Figure 4, right panel) using the *circlize* software library in R, restricting the visualization to a subset of networks of interest.

### 10. Identifying Highly Varying Network Pairs

We quantified variation of pairs of networks across 15 connectomic neuromarkers (4 for the factor model and 11 for individual tasks). For each neuromarker, we summed suprathreshold connections for each pair of networks separately for positive and negative connections, yielding 240 values per neuromarker. We next calculated the variance of these values across the 15 neuromarkers. We selected the 21 most varying network pairs (which together accounted for over 50% of the variance across the 15 neuromarkers) and displayed the number of suprathreshold connections for each network pair for each neuromarker with a bubble heat map.

### 11. Data Availability

The ABCD data used in this report came from NDA Study 721, 10.15154/1504041, which can be found at https://nda.nih.gov/study.html?id=721.

## ACKNOWLEDGEMENTS

Data used in the preparation of this article were obtained from the Adolescent Brain Cognitive Development (ABCD) Study (https://abcdstudy.org), held in the NIMH Data Archive (NDA). This is a multisite, longitudinal study designed to recruit more than 10,000 children age 9-10 and follow them over 10 years into early adulthood. The ABCD Study is supported by the National Institutes of Health and additional federal partners under award numbers U01DA041022, U01DA041028, U01DA041048, U01DA041089, U01DA041106, U01DA041117, U01DA041120, U01DA041134, U01DA041148, U01DA041156, U01DA041174, U24DA041123, and U24DA041147. A full list of supporters is available at https://abcdstudy.org/nih-collaborators. A listing of participating sites and a complete listing of the study investigators can be found at _. ABCD consortium investigators designed and implemented the study and/or provided data but did not necessarily participate in analysis or writing of this report. This manuscript reflects the views of the authors and may not reflect the opinions or views of the NIH or ABCD consortium investigators. The ABCD data repository grows and changes over time. The ABCD data used in this report came from NDA Study 721, 10.15154/1504041, which can be found at https://nda.nih.gov/study.html?id=721.

This work was supported by the following grants from the United States National Institutes of Health, the National Institute on Drug Abuse, and the National Institute on Alcohol Abuse and Alcoholism: R01MH107741 (CS), U01DA041106 (CS, MH, LH), T32 AA007477 (DC, AW). In addition, CS was supported by a grant from the Dana Foundation David Mahoney Neuroimaging Program. This research was supported in part through computational resources and services provided by Advanced Research Computing at the University of Michigan, Ann Arbor.

## AUTHOR CONTRIBUTIONS

Conceptualization: CS, MA, AT, SR, MH; Methodology: CS, DC, MA, AT, SR; Formal Analysis: CS, DC, MA, AT, SR; Data Curation: MA, SR, TG; Writing – Original Draft: CS; Writing – Reviewing and Editing; AW, CS, LH, MA, MH, SR, TG; Visualization: MA, AT, SR; Supervision: CS, MH; Funding Acquisition: CS, MH.

## REFERENCES

1. Rosenberg, M. D. et al. A neuromarker of sustained attention from whole-brain functional connectivity. Nat. Neurosci. 19, 165–171 (2016).

2. Greene, A. S., Gao, S., Scheinost, D. & Constable, R. T. Task-induced brain state manipulation improves prediction of individual traits. Nature communications 9, 1–13 (2018).

3. Dubois, J., Galdi, P., Paul, L. K. & Adolphs, R. A distributed brain network predicts general intelligence from resting-state human neuroimaging data. Philos Trans R Soc Lond B Biol Sci 373, (2018).

4. Kessler, D., Angstadt, M. & Sripada, C. Brain Network Growth Charting and the Identification of Attention Impairment in Youth. JAMA Psychiatry 73, 481–489 (2016).

5. Sripada, C. et al. Prediction of Neurocognition in Youth From Resting State fMRI. Molecular Psychiatry 1–19 (2019) doi:10.1101/495267.

6. Sui, J., Jiang, R., Bustillo, J. & Calhoun, V. Neuroimaging-based Individualized Prediction of Cognition and Behavior for Mental Disorders and Health: Methods and Promises. Biological Psychiatry (2020).

7. Diamond, A. Executive functions. Annual review of psychology 64, 135–168 (2013).

8. Casey, B. J., Tottenham, N. & Fossella, J. Clinical, imaging, lesion, and genetic approaches toward a model of cognitive control. Dev. Psychobiol. 40, 237–254 (2002).

9. Banich, M. T. Executive function: The search for an integrated account. Current directions in psychological science 18, 89–94 (2009).

10. Luerssen, A. & Ayduk, O. Executive functions promote well-being: Outcomes and mediators. in The happy mind: Cognitive contributions to well-being 59–75 (Springer, 2017).

11. Gottfredson, L. S. Why g matters: The complexity of everyday life. Intelligence 24, 79–132 (1997).

12. Snyder, H. R., Miyake, A. & Hankin, B. L. Advancing understanding of executive function impairments and psychopathology: bridging the gap between clinical and cognitive approaches. Frontiers in psychology 6, 328 (2015).

13. McTeague, L. M., Goodkind, M. S. & Etkin, A. Transdiagnostic impairment of cognitive control in mental illness. Journal of psychiatric research 83, 37–46 (2016).

14. Pennington, B. F. & Ozonoff, S. Executive functions and developmental psychopathology. Journal of child psychology and psychiatry 37, 51–87 (1996).

15. Banich, M. T. et al. Cognitive control mechanisms, emotion and memory: a neural perspective with implications for psychopathology. Neuroscience & Biobehavioral Reviews 33, 613–630 (2009).

16. Sripada, C. & Weigard, A. S. Impaired Evidence Accumulation as a Transdiagnostic Vulnerability Factor in Psychopathology. (2020).

17. Niendam, T. A. et al. Meta-analytic evidence for a superordinate cognitive control network subserving diverse executive functions. Cogn Affect Behav Neurosci 12, 241–268 (2012).

18. Cole, M. W. & Schneider, W. The cognitive control network: Integrated cortical regions with dissociable functions. NeuroImage 37, 343–360 (2007).

19. Duncan, J. & Owen, A. M. Common regions of the human frontal lobe recruited by diverse cognitive demands. Trends in Neurosciences 23, 475–483 (2000).

20. Gray, J. R., Chabris, C. F. & Braver, T. S. Neural mechanisms of general fluid intelligence. Nat. Neurosci. 6, 316–322 (2003).

21. Bassett, D. S. & Sporns, O. Network neuroscience. Nature neuroscience 20, 353 (2017).

22. Sporns, O. Contributions and challenges for network models in cognitive neuroscience. Nat. Neurosci. 17, 652–660 (2014).

23. Smith, S. M. et al. Functional connectomics from resting-state fMRI. Trends in Cognitive Sciences 17, 666–682 (2013).

24. Woo, C.-W., Chang, L. J., Lindquist, M. A. & Wager, T. D. Building better biomarkers: brain models in translational neuroimaging. Nature Neuroscience 20, 365–377 (2017).

25. Spearman, C. ‘ General Intelligence,’ objectively determined and measured. The American Journal of Psychology 15, 201–292 (1904).

26. Carroll, J. B. Human cognitive abilities: A survey of factor-analytic studies. (Cambridge University Press, 1993).

27. Karr, J. E. et al. The unity and diversity of executive functions: A systematic review and reanalysis of latent variable studies. Psychological bulletin 144, 1147 (2018).

28. Bloemen, A. J. P. et al. The association between executive functioning and psychopathology: general or specific? Psychological medicine 48, 1787–1794 (2018).

29. Horn, J. L. Organization of abilities and the development of intelligence. Psychological review 75, 242 (1968).

30. Carroll, J. B. The higher-stratum structure of cognitive abilities: Current evidence supports g and about ten broad factors. in The scientific study of general intelligence 5–21 (Elsevier, 2003).

31. Volkow, N. D. et al. The conception of the ABCD study: From substance use to a broad NIH collaboration. Developmental cognitive neuroscience 32, 4–7 (2018).

32. Karcher, N. R. & Barch, D. M. The ABCD study: understanding the development of risk for mental and physical health outcomes. Neuropsychopharmacology 1–13 (2020).

33. Luciana, M. et al. Adolescent neurocognitive development and impacts of substance use: overview of the adolescent brain cognitive development (ABCD) baseline neurocognition battery. Developmental cognitive neuroscience (2018).

34. Zinbarg, R. E., Revelle, W., Yovel, I. & Li, W. Cronbach’s α, Revelle’s β, and McDonald’s ω H: Their relations with each other and two alternative conceptualizations of reliability. psychometrika 70, 123–133 (2005).

35. Sripada, C. et al. Basic Units of Inter-Individual Variation in Resting State Connectomes. Scientific Reports 9, 1900 (2019).

36. Sripada, C., Angstadt, M., Rutherford, S., Taxali, A. & Shedden, K. Toward a “treadmill test” for cognition: Improved prediction of general cognitive ability from the task activated brain. Human Brain Mapping (2020).

37. Scheinost, D. et al. Ten simple rules for predictive modeling of individual differences in neuroimaging. NeuroImage (2019).

38. Casey, B. J. et al. The adolescent brain cognitive development (ABCD) study: imaging acquisition across 21 sites. Developmental cognitive neuroscience (2018).

39. Marek, S. et al. Towards Reproducible Brain-Wide Association Studies. bioRxiv (2020).

40. Finn, E. S. et al. Functional connectome fingerprinting: identifying individuals using patterns of brain connectivity. Nature Neuroscience 18, 1664–1671 (2015).

41. Duncan, J. et al. A neural basis for general intelligence. Science 289, 457–460 (2000).

42. Luna, B., Marek, S., Larsen, B., Tervo-Clemmens, B. & Chahal, R. An integrative model of the maturation of cognitive control. Annual review of neuroscience 38, 151–170 (2015).

43. Casey, B. J., Tottenham, N., Liston, C. & Durston, S. Imaging the developing brain: what have we learned about cognitive development? Trends in Cognitive Sciences 9, 104–110 (2005).

44. Durston, S. & Casey, B. J. What have we learned about cognitive development from neuroimaging? Neuropsychologia 44, 2149–2157 (2006).

45. Cole, M. W. et al. Multi-task connectivity reveals flexible hubs for adaptive task control. Nature neuroscience 16, 1348 (2013).

46. Gratton, C., Sun, H. & Petersen, S. E. Control networks and hubs. Psychophysiology 55, e13032 (2018).

47. Cui, Z. et al. Individual Variation in Functional Topography of Association Networks in Youth. Neuron (2020).

48. Fair, D. A. et al. Development of distinct control networks through segregation and integration. Proc. Natl. Acad. Sci. U.S.A. 104, 13507–13512 (2007).

49. Barber, A. D., Caffo, B. S., Pekar, J. J. & Mostofsky, S. H. Developmental changes in within- and between-network connectivity between late childhood and adulthood. Neuropsychologia 51, 156–167 (2013).

50. Grayson, D. S. & Fair, D. A. Development of large-scale functional networks from birth to adulthood: A guide to the neuroimaging literature. NeuroImage 160, 15–31 (2017).

51. Di Martino, A. et al. Unraveling the Miswired Connectome: A Developmental Perspective. Neuron 83, 1335–1353 (2014).

52. Paus, T. Mapping brain maturation and cognitive development during adolescence. Trends in Cognitive Sciences 9, 60–68 (2005).

53. Green, M. F. What are the functional consequences of neurocognitive deficits in schizophrenia? The American journal of psychiatry (1996).

54. Heinrichs, R. W. & Zakzanis, K. K. Neurocognitive deficit in schizophrenia: a quantitative review of the evidence. Neuropsychology 12, 426 (1998).

55. Mesholam-Gately, R. I., Giuliano, A. J., Goff, K. P., Faraone, S. V. & Seidman, L. J. Neurocognition in first-episode schizophrenia: a meta-analytic review. Neuropsychology 23, 315 (2009).

56. Barkley, R. A. Behavioral inhibition, sustained attention, and executive functions: Constructing a unifying theory of ADHD. Psychological Bulletin 121, 65–94 (1997).

57. Willcutt, E. G., Doyle, A. E., Nigg, J. T., Faraone, S. V. & Pennington, B. F. Validity of the executive function theory of attention-deficit/hyperactivity disorder: a meta-analytic review. Biol. Psychiatry 57, 1336–1346 (2005).

58. Bates, M. E., Bowden, S. C. & Barry, D. Neurocognitive impairment associated with alcohol use disorders: implications for treatment. Experimental and clinical psychopharmacology 10, 193 (2002).

59. Stevens, L. et al. Impulsivity as a vulnerability factor for poor addiction treatment outcomes: a review of neurocognitive findings among individuals with substance use disorders. Journal of substance abuse treatment 47, 58–72 (2014).

60. Snyder, H. R. Major depressive disorder is associated with broad impairments on neuropsychological measures of executive function: a meta-analysis and review. Psychological bulletin 139, 81 (2013).

61. Fossati, P., Ergis, A. M. & Allilaire, J. F. Executive functioning in unipolar depression: a review. L’encéphale 28, 97–107 (2002).

62. Brislin, S. et al. Differentiated Nomological Networks of Internalizing, Externalizing, and the General Factor of Psychopathology (“P factor”) in Emerging Adolescence in the ABCD study. (2020).

63. Lahey, B. B. et al. Is there a general factor of prevalent psychopathology during adulthood? Journal of abnormal psychology 121, 971 (2012).

64. Caspi, A. & Moffitt, T. E. All for one and one for all: Mental disorders in one dimension. American Journal of Psychiatry 175, 831–844 (2018).

65. Smith, G. T., Atkinson, E. A., Davis, H. A., Riley, E. N. & Oltmanns, J. R. The general factor of psychopathology. Annual Review of Clinical Psychology 16, 75–98 (2020).

66. Garavan, H. et al. Recruiting the ABCD sample: design considerations and procedures. Developmental cognitive neuroscience 32, 16–22 (2018).

67. Barch, D. M. et al. Demographic, physical and mental health assessments in the adolescent brain and cognitive development study: Rationale and description. Developmental cognitive neuroscience (2017).

68. Hagler, D. J. et al. Image processing and analysis methods for the Adolescent Brain Cognitive Development Study. bioRxiv 457739 (2018) doi:10.1101/457739.

69. Esteban, O. et al. fMRIPrep: a robust preprocessing pipeline for functional MRI. Nature methods 16, 111 (2019).

70. Gorgolewski, K. et al. Nipype: a flexible, lightweight and extensible neuroimaging data processing framework in python. Frontiers in neuroinformatics 5, 13 (2011).

71. Gordon, E. M. et al. Generation and Evaluation of a Cortical Area Parcellation from Resting-State Correlations. Cereb. Cortex 26, 288–303 (2016).

72. Tian, Y., Margulies, D. S., Breakspear, M. & Zalesky, A. Hierarchical organization of the human subcortex unveiled with functional connectivity gradients. bioRxiv (2020).

73. Diedrichsen, J. et al. Imaging the deep cerebellar nuclei: a probabilistic atlas and normalization procedure. Neuroimage 54, 1786–1794 (2011).

74. Lindquist, M. A., Geuter, S., Wager, T. D. & Caffo, B. S. Modular preprocessing pipelines can reintroduce artifacts into fMRI data. Human brain mapping 40, 2358–2376 (2019).

